# The molecular core of transcriptome responses to abiotic stress in plants: a machine learning-driven meta-analysis

**DOI:** 10.1101/2024.01.24.576978

**Authors:** Raul Sanchez-Munoz, Thomas Depaepe, Marketa Samalova, Jan Hejatko, Isiah Zaplana, Dominique Van Der Straeten

## Abstract

Understanding how plants adapt their physiology to overcome severe stress conditions is vital in light of the current climate crisis. This remains a challenge given the complex nature of the underlying molecular mechanisms. To provide a full picture of stress mitigation mechanisms, an exhaustive analysis of publicly available stress-related transcriptomic data was conducted. We combined a meta-analysis with an unsupervised machine learning algorithm to identify a core of stress-related genes. To ensure robustness and biological significance of the output, often lacking in meta-analyses, a three-layered biovalidation was incorporated. Our results present a ‘stress gene core’, a set of key genes involved in plant tolerance to a multitude of adverse environmental conditions rather than specific ones. In addition, we provide a biologically validated database to assist in design of multi-stress resilience. Taken together, our results pave the way towards future-proof sustainable agriculture.

**Teaser:** Using a machine learning-driven meta-analysis, a plant ‘stress gene core’ was identified as a hub mediating multi-stress regulation

## Introduction

Environmental factors such as light, temperature, and water availability are key signals steering plant growth and development. The current climate crisis generated by global warming steadily increases the number of geographical regions suffering from extreme environmental conditions (1). The latter create situations in which plants, even if their survival is not severely compromised, grow under suboptimal conditions, which ultimately hamper their growth and reduce yield (2). Considering that the vast majority of the world’s arable surface is exposed to biotic and abiotic stress conditions – with estimations of up to 30% and 82% of worldwide crop productivity decay, respectively – and that climate change hits the planet faster than anticipated (IPCC report, 2023), we face an imminent global agronomical crisis (3,4). Nevertheless, plants have evolved sophisticated mechanisms to cope with adverse environmental conditions.

Several efforts have been focused on unravelling stress mitigation pathways with the aim of engineering novel strategies for the improvement of plant stress tolerance, though with limited success (5). This could be caused by the diversity in stress responses, as well as the complexity of each stress type and its associated signaling pathway. In addition, environmental conditions usually impose multifactorial stimuli rather than a juxtaposition of individual stresses, which plants perceive as a combination of different inputs (6). Thus, plants require a strict and coordinated communication between different tissues and organs to optimally respond to unfavorable conditions (7). Systemic responses between distant tissues or organs have been demonstrated by the application of localized stress stimuli, confirming the existence of efficient communication routes within plants (8).

Besides these systemic and coordinated responses, different tissues also perform specific tasks, related to their primary physiological functions (9). Therefore, inter-organ communication and tissue-specific responses are crucial for optimal stress mitigation. In addition, responses to stress also depend on its duration. At early stages, stress signaling is focused on rapid biochemical, biophysical, molecular and physiological alterations that avoid and/or reduce irreversible cellular damage (10). As stress persists, the acclimation process starts, modulating growth and development to ensure survival and, concomitantly regulating the mechanisms initiated during the early stages (11-13). The gaseous hormone ethylene (ETH) is one of the main players in environmental adaptation (14). Transcriptional analyses have detected several groups of important stress-related transcription factors (TF) that are directly influenced by ETH, such as AP2/ERF (Apetala 2/Ethylene responsive factor), NAC (NAM (No Apical Meristem), ATAF1/2 (Activating Factor 1/2), CUC2 (Cup-shaped Cotyledon 2)) and WRKYs, among others (15). Nevertheless, many aspects of ETH-mediated stress adaptation are still poorly understood, often clouded by the intricate crosstalk with other regulatory pathways.

Although stress responses are generally defined as highly specific and sometimes potentially antagonizing (16), several clues evidence the existence of a shared molecular core acting as a signaling hub to coordinate multiple stress stimuli (17). This is further supported by the finding that stress priming – the exposure to mild stress conditions in order to develop posterior stress tolerance – can induce cross-tolerance (18). Heat and cold cross-tolerance is a common and well-known example due to their physiological similitudes (19), but this relationship has also been detected between heat and cadmium (20), or cold, salt, and drought (21). Together, this points towards the existence of a ‘stress gene core’, responsible for the coordination of specific responses towards physiological adaptation upon compound stresses, though the exact players remain to be determined.

However, several factors impede the identification of such stress core, predominantly related to the enormous complexity of the network controlling plant stress tolerance and the multifactorial nature of stress. A direct consequence of this complexity is the fact that the modification of specific elements to enhance stress tolerance can generate unpredictable and unwanted consequences (22). Therefore, a deeper understanding of factors shared between stress signaling routes, both in spatial and temporal contexts, is vital. Moreover, the large quantity of potential genes involved in stress responses infers that the experimental efforts needed to understand all the players involved are overwhelming. Hence, it is clear that robust *in silico* methods are crucial to gain insights into the systems’ complexity and guide experimental confirmation.

The extensive study of transcriptional changes by means of next-generation sequencing methods provides a rich and diverse library of data. Nevertheless, single transcriptomic analyses can produce contradictory conclusions, driven by experimental differences such as the type of treatment, the severity of stress stimuli, the time range of treatment, tissue and plant age, and/or sample size (23), or lead to biased, misleading results. The combination of multiple transcriptomic data with meta-analysis approaches has been proposed as a method to bypass these limitations (24), providing solid input for the definition of a gene core. However, the design and implementation of meta-analyses are not trivial, since they require the combination of powerful statistical methods without losing biological significance (25).

For that reason, we have performed a meta-analysis combining all publicly available stress-related *Arabidopsis thaliana* transcriptomic data, after careful and individual consideration of the suitability of each transcriptome in order to ensure biological significance of the analysis. Our aim was to identify an abiotic stress gene core, given the impact of abiotic stress on crop yield (4) and exacerbation of stress by global climate change. Such complex datasets require, in addition to a reliable meta-analysis method, a potent data mining tool to extract valuable information. Recently, the high potential of machine learning techniques for data analysis has been extensively demonstrated, as well as the limitations and flaws that need to be taken into consideration for its proper usage and interpretation (26).

The high-dimensional dataset derived from the combination of all single transcriptomes requires an efficient machine learning method able to cope with such vaste data. Support Vector Machine (SVM) is easy to implement for classification of complex multi-dimensional datasets (27). In particular, an unsupervised version of the standard SVM, called SVM Clustering, was selected for this work, as it preserves all the key properties of a standard SVM while, concurrently, avoiding the limitations of purely supervised methods, e.g. overfitting (28-30). In order to properly control the reproducibility and robustness of our methodology, and to increase the biological significance of the results, often lacking in meta-analyses, we have designed a three-layered biovalidation, including experimental validation of some key stress-related genes detected in our analysis.

This work presents the first machine learning-driven meta-analysis of abiotic stress-related plant genes including all publicly available Arabidopsis datasets. The final output is a list of genes forming the plant ‘abiotic stress gene core’. Rather than being stress-specific – as when derived from single transcriptomic analyses – these genes represent potential hubs of general stress responses in plants. Therefore, this core has the potential to be a ‘gold mine’ for the development of multi-resistant crop varieties. Secondly, the analysis offers trustworthy information on the temporal and spatial expression of key stress genes and regulatory gene families. Thirdly, novel insights are gained into the presently unknown functionality of relevant genes, as well as into key members within large multigene families. Overall, both our approach and the obtained results represent an important step forward in the field of plant systems biology, offering a powerful methodology to identify biologically relevant core genes, supporting more robust engineering strategies for future development of stress-resistant plants.

## Results

### Construction of DEG libraries and hierarchical clustering

After screening and filtering all available abiotic stress-related transcriptomic datasets from the Gene Expression Omnibus (GEO) database, 500 individual transcriptomes were analyzed for differentially expressed genes (DEGs) (Fig. S1, Suppl. File 1). Based on the kinetics of individual transcriptomic analyses, the lists of DEGs were combined in early (from 1 to 6 hours) and late responses (from 12 to 24 hours) (Suppl. File 2; for more details, see Materials and Methods). Prior to further analyses, a first biovalidation step was performed. To confirm the suitability and biological relevance of each DEG library, 6 marker genes were selected per stress condition based on their empirically determined expression. Subsequently, the presence of these markers was assessed for each stress, taking into account temporal and spatial specificity (Suppl. Table 1). All passed the first biological validation test with at least 5 out of 6 markers, demonstrating both the suitability and the accuracy of the DEG libraries.

The number of DEGs obtained in each stress, tissue, and timepoint combination provided a first insight into the regulation of abiotic stress responses (Fig. 1A and 1D). In general, the responses were balanced considering up- and down-regulation and number of DEGs between tissues and timepoints. A Hierarchical Clustering Analysis (HCA) grouped certain stresses by DEG modules, suggesting physiological resemblances between them (e.g. salt and osmotic stress in roots and shoots; partial and complete submergence in roots; wounding and drought in shoots; and early exposure to UV and high light in shoots; Fig. 1B-C, 1E-F).

**Figure 1.**
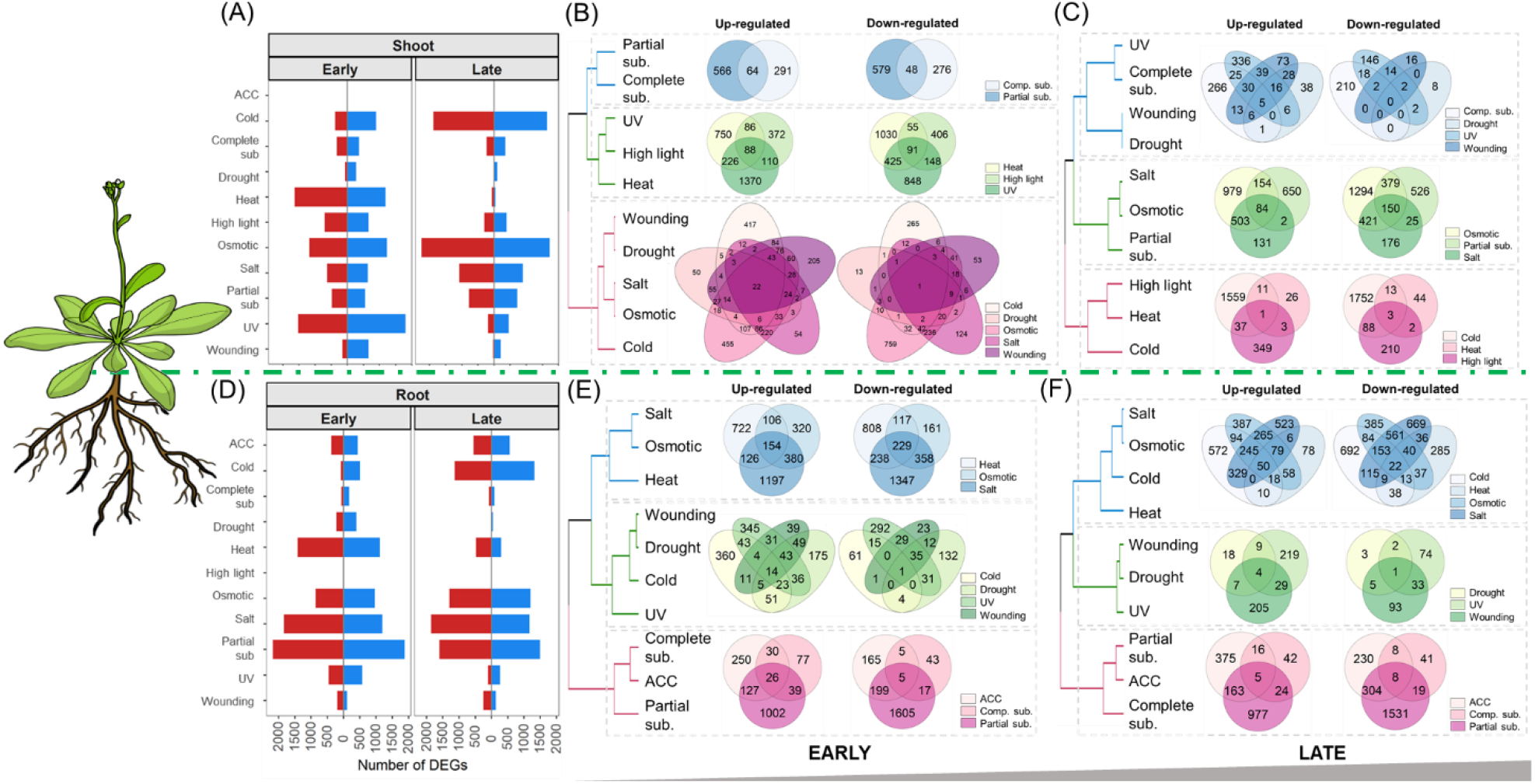
Distribution of differentially expressed genes (DEGs) per stress, tissue, and time point. **(A)** Number of DEGs detected in each of the studied conditions in shoot tissue for early and late time points. Upregulated DEGs are colored in blue, downregulated in red. **(B and C)** Hierarchical Clustering Analysis (HCA) of the different stress conditions by DEG modules in shoots for early **(B)** and late **(C)** time points. **(D)** Number of DEGs detected in each of the studied conditions in root tissue for early and late timepoints. Upregulated DEGs are colored in blue, downregulated in red. **(E and F)** HCA of the different stress conditions by DEG modules in roots for early **(E)** and late **(F)** time points. Branch length is based on cluster distance.

In addition, the comparison of DEG modules corresponding to early and late time points highlighted the existence of a set of transcriptional responses in roots that are relatively stable in time. Regarding tissue specificity, responses in roots and shoots exhibited some overlap in the early timepoints, though clearly diverged over time (Fig. 1; Fig. S2-3). This advocates the existence of a tissue-specific mechanism to respond to stress conditions, especially when maintained over time (for more details, see Suppl. Text).

### Support Vector Machine (SVM) Clustering based classification for stress core determination

In order to identify a set of central actors in stress signaling, i.e. as part of a ‘stress gene core’, we have computed the meta-*p*-values for all studied genes (for details, see Material and Methods). We took into account their transcriptional changes in all stress conditions in four different datasets: roots early, roots late, shoots early and shoots late, and used the SVM Clustering (an unsupervised version of the standard SVM). First, a frequency-based pre-classification was performed. The genes appearing as DEGs in at least 5 of the studied conditions were assigned to the positive class (class 1), while the rest was assigned to the negative class (class 0). Subsequently, the meta-*p*-value dataset containing the information of all 10 stress conditions for the complete set of around 12,000 genes, was re-classified using the SVM Clustering algorithm (for details, see Material and Methods section). This analysis classifies genes depending on their distribution in the 10-dimensional space taking into account the distribution of meta-*p-*values (reflecting how significant the expression changes are under each condition).

SVM Clustering categorized the vast majority of genes as not relevant (class 0; approximately 99%), coinciding with the frequency-based pre-classification (Fig. 2A). Around 5 to 30% of the genes pre-classified as relevant (32, 6, 82 and 14 genes for the early-root, late-root, early-shoot and late-shoot responses, respectively) were refuted after SVM Clustering (1>0). In contrast, a few genes pre-classified as not relevant (0) were included in the final SVM gene core (0>1) (2, 4, 7 and 0 genes from the early-root, late-root, early-shoot and late-shoot responses, respectively). The significant number of genes pre-classified as relevant but considered irrelevant by the SVM Clustering highlights the discriminatory power of the SVM Clustering-based classification, refining the pre-classification established on the distribution of the complete set of meta-*p*-values. Based on this, four sets of core genes, coined SVM gene cores, with significant transcriptional alterations in all the studied conditions, were identified: 118 genes for early responses in roots, 108 for late responses in roots, 185 for early responses in shoots and 74 for late responses in shoots (Fig. 2A).

**Figure 2.**
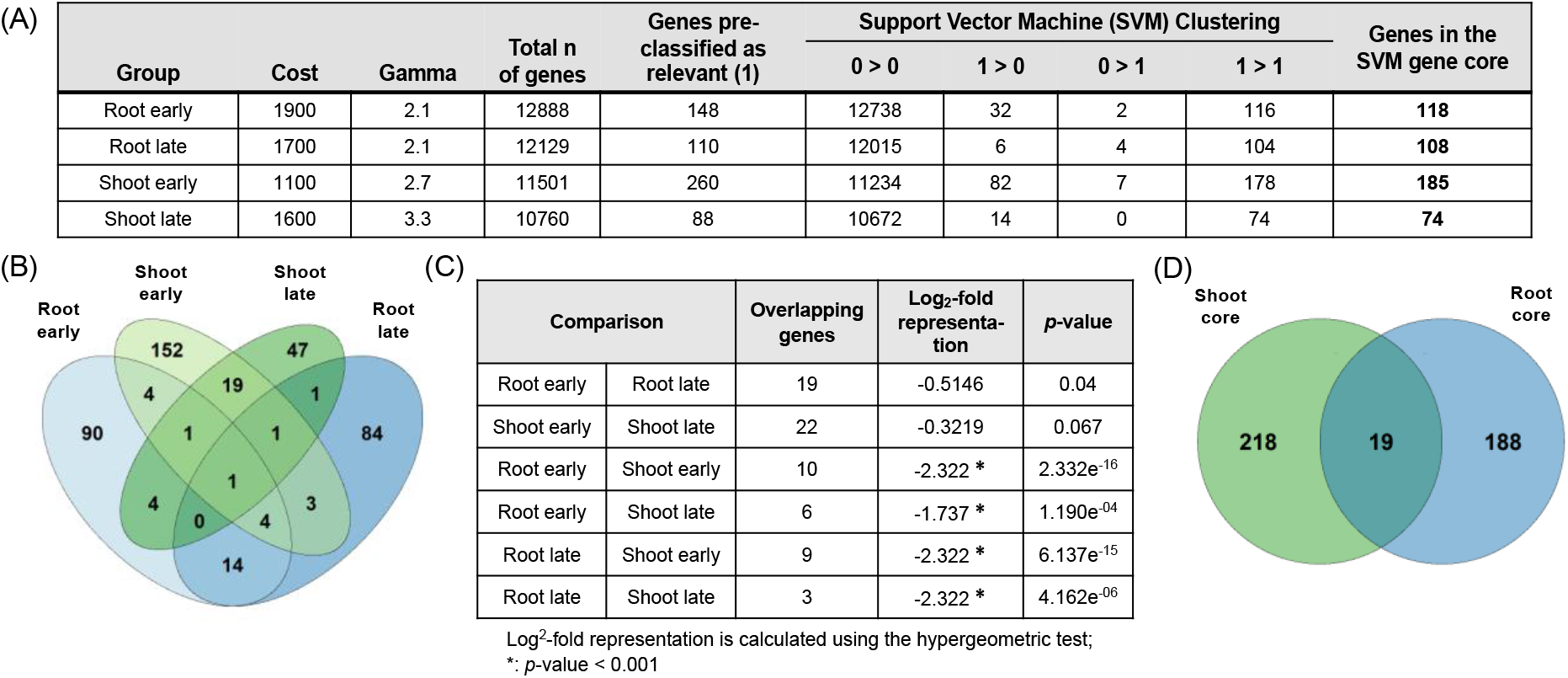
Support Vector Machine (SVM) Clustering for the determination of the stress gene cores. **(A)** SVM-specific parameters (cost and gamma) used for the SVM Clustering of each tissue and timepoint dataset. The number of genes forming each SVM gene core is highlighted in bold. A pre-classification was performed first, where relevant genes appearing as a DEG in at least 5 of the stress conditions studied were assigned to the positive class (class 1). The results of the SVM Clustering are defined as 0>0 (genes marked as not relevant and stated as such by the algorithm), 1>0 (genes marked as relevant but not stated as such by the algorithm), 0>1 (genes not marked as relevant but stated as such by the algorithm) and 1>1 (genes marked as relevant and stated as such by the algorithm). **(B)** Venn diagram representing the overlap between different SVM gene cores. **(C)** Overlaps between the SVM gene cores. The value, positive (overrepresentation) or negative (underrepresentation), and the significance of the overlaps are indicated in the table. A hypergeometric test was used to detect the significance of the overlaps, considering a *p*-value < 0.001 as significant due to the reduced number of genes in the comparisons. **(D)** Venn diagram representing the overlapp between the root and shoot gene cores.

Interestingly, the comparison between tissues (root versus shoot in all timepoint combinations; Fig. 2B-C), as opposed to comparisons between timepoints within a tissue (root early versus root late; shoot early versus shoot late), revealed a significant underrepresentation of overlapping genes. Therefore, considering gene response composition, tissue-specificity is stronger than temporal specificity, supporting the results obtained by the qualitative DEG classification (Fig. 1, Fig. S3). For that reason, and due to the non-significant differences between timepoints (Fig. 2C), the genes belonging to the SVM gene cores per tissue were combined, resulting in a final dataset of 207 genes forming the root gene core and 237 genes forming the shoot gene core after removal of duplicates (Fig. 2D). Despite the tissue-specificity of the SVM gene cores, a set of 19 genes is shared between the root and shoot gene cores, which could encompass fundamental proteins with tissue-independent functions. These predominantly cover genes involved in cell wall maintenance and membrane integrity *(EXPANSIN A1 (EXPA1), LIPID TRANSFER PROTEIN 2 (LTP2),* dehydrins such as *COLD-REGULATED 47 (COR47)* and *LOW TEMPERATURE-INDUCED 30 (LTI30),* and *BLUE COPPER BINDING PROTEIN (BCB)) (*31-33), in addition to some uncharacterized genes (e.g. *AT1G19380* and *AT5G19875*). The complete list of genes that form the different SVM gene cores as well as their overlap can be found in the Suppl. File 3.

We studied the composition of the gene cores in terms of annotated biological functions (Gene Ontology (GO) enrichment analysis) and gene families. While the most significant function enriched in the shoot core was the response to water (GO:0009415), response to hypoxia appeared to be major one in roots (GO:0071456) (Fig. S2, Fig. S4). This reflects that stressed shoots prioritize on maintenance of water homeostasis, while roots mostly try to maintain normoxia. In addition, amino acid transporters were enriched in the shoot core, while EXPANSINs, related to cell wall remodeling, appeared on the forefront in the root core (for a complete GO and gene family analysis, see Suppl. Text).

### Protein networks related to the SVM gene cores

To further elucidate the functionality of shoot and root cores, we constructed a protein network representing both physical and functional interactions. Subsequently, a *k*-means clustering method was applied to obtain protein clusters based on known interactions in order to provide further evidence of their biological roles (for details, see Material and Methods).

For shoots, four clusters were identified (Fig. 3A). The blue and red clusters contained the largest number of proteins (28%). However, given its higher degree of connectivity, the blue cluster was considered to be a key cluster within the shoot core (Fig. 3A-B). In order to support the biological relevance of the different clusters, we performed a biological processes GO enrichment for each one individually (Fig. 3C-F). As expected, the biological responses of the blue cluster largely overlapped with those of the overall shoot core (Fig. S4), with ‘response to water deprivation’ (GO:0009414) as the most significant GO term (Fig. 3F). Three WRKY TFs appeared to act as central nodes in the interaction network, strongly interacting among themselves (WRKY33, WRKY46 and WRKY18) (Fig. 3B). In addition, MAP KINASE KINASE 9 (MKK9), a Mitogen Activated Protein Kinase (MAPK) protein, directly interacts with WRKY33, the central protein in the interactome, suggesting an important regulatory role for MKK9 as well. Furthermore, the interaction network also highlighted other previously discovered stress genes as part of the stress signaling core, including the mitochondrial *ALTERNATIVE OXIDASE 1A* (*AOX1a*), as well as 31 unannotated genes, hence uncovering their function (Suppl. File 4).

**Figure 3.**
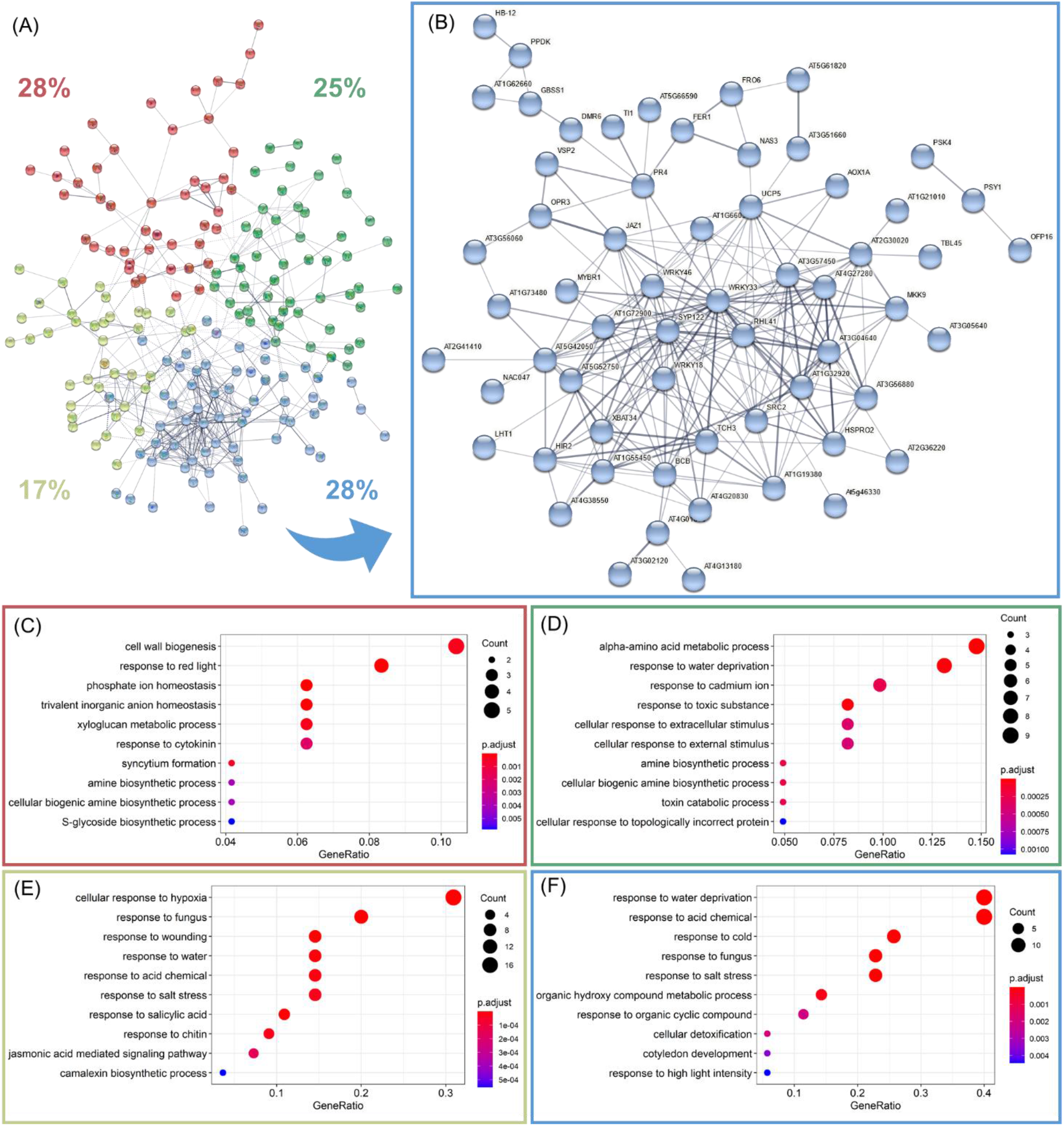
Protein interaction network based on the shoot gene core and GO enrichment of each cluster. **(A)** Protein interaction network representing functional and physical interactions between proteins of the shoot gene core. Node colors represent different clusters depending on neighbor interactions;connecting lines thickness represents the strength of the connections based on experimental data. Percentages indicate the number of proteins forming the cluster in relation to the total number of genes of the SVM gene core. **(B)** Cluster that contains the highest number of proteins of the 4 clusters. The name or AGI number of each gene is indicated next to the corresponding node. **(C to F)**. GO terms related to biological processes obtained during the GO enrichment of each cluster. The colored square indicates which GO analysis corresponds with each cluster highlighted in (A). The parameter p.adjust represents the significance of the group (*p*-value < 0.05 was used as cutoff). Count indicates the number of genes inside the category. GeneRatio reflects the percentage of DEGs in the complete GO category.

The red cluster contained proteins that are mainly related to the maintenance of the cell wall integrity (GO:0042546, GO:0010411, GO:0006949) (Fig. 3C). The green cluster was marked by ‘alpha-amino acid metabolic process’ (GO:1901605) and ‘response to water deprivation’ (GO:0009414) as main GO terms, reflecting its role in both metabolism and responses to water availability (Fig. 3D). Lastly, the smallest cluster (yellow) contained proteins involved in hypoxia responses (GO:0071456), together with proteins related to other stress responses (biotic and wounding stress) (Fig. 3E). In conclusion, it is evident that shoot stress signaling mainly mitigates alterations in water status and conserves cellular water homeostasis. In addition, MKK9 and WRKY TFs, specifically WRKY33, appear to be pivotal in the regulation of these responses.

A similar topology was obtained for the root core interaction network (Fig. 4A). Of the four clusters, two contained the maximum number of proteins (representing 28% of the total number in the core) and, one of them (colored in blue), exhibited the highest number of connections within the network (Fig. 4B). As expected, the blue cluster showed the GO category that characterized the root core (GO:0071456: ‘cellular response to hypoxia’). The second most relevant GO term (‘secondary metabolic process’ (GO:0019748)) covered genes involved in lignan biosynthesis, such as *BCB*, and phenylpropanoid biosynthesis (*KISS ME DEADLY 1* and *4*; *KMD1*/*4*). In addition, defense responses seemed to play an important role in this blue cluster (GO:0031347), as well as responses to external stimuli (GO:0009605) and to ethylene (GO:0009723), indicating a relevant role in environmental interactions (Fig. 4F). The MAPK protein MPK11 was found at a central position in the interaction network, possibly coordinating the activity of the remaining members of the blue cluster. Interestingly, the ETH receptor ETHYLENE RESPONSE 2 (ETR2) and the downstream TF ETHYLENE RESPONSE FACTOR 2 (ERF2), which is induced by ETH (34), were also present in this cluster, corroborating a pivotal role of ETH in the generic stress response, at least in roots.

**Figure 4.**
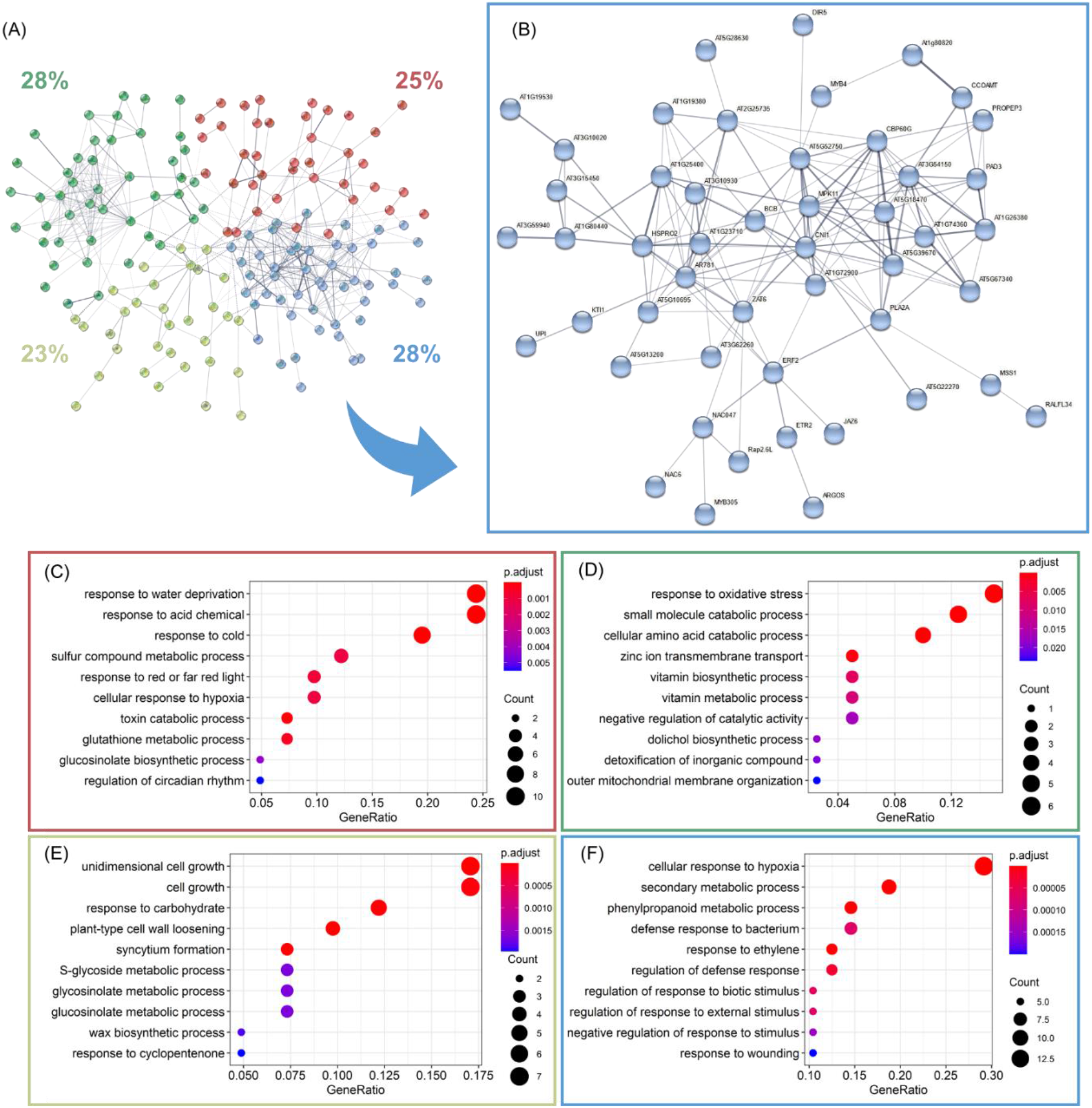
Protein interaction network of the root gene core and GO enrichment of each cluster. **(A)** Protein interaction network representing functional and physical interactions between proteins forming the root gene core. Node colors represent different clusters depending on neighbor interactions, and connecting lines thickness represents the strength of the connections based on experimental data. Percentages indicate the number of proteins forming the cluster in relation to the total of genes of the SVM gene core. **(B)** Cluster that contains the highest number of proteins of the 4 clusters. The name or AGI number of each gene is indicated next to the corresponding node. **(C to F)**. GO terms related to biological processes obtained during the GO enrichment for each cluster. The colored square indicates which GO analysis corresponds with each cluster highlighted in (A). The parameter p.adjust represents the significance of the group (*p*-value < 0.05 was used as cutoff). Count indicates the number of genes inside the category. GeneRatio reflects the percentage of DEGs in the complete GO category.

The green cluster (Fig. 4D) was characterized by GO terms related to oxidative stress responses (GO:0006979), cellular metabolism of amino acids (GO:0009063) and vitamins (GO:0009110; GO:0006766), and transport of inorganic compounds (GO:0006829), revealing a potential role for the maintenance of shoot metabolism and physiology in root responses. Finally, both red and yellow clusters showed a reduced number of proteins and interaction levels compared to the previous ones (Fig. 4C and 4E). Interestingly, the yellow cluster showed GO terms involved mainly in cell wall homeostasis and modification (GO:0009826; GO:0016049; GO:0009828; GO:0006949; GO:0010025) while the red one included GO terms water responses (GO:0009415), and hypoxia (GO:0071456), among others.

Overall, we conclude that shoot stress responses are mostly related to the maintenance of water potential and homeostasis and, secondary, to the maintenance of normoxia levels; while in roots, the opposite trend is observed. In addition, growth regulation and metabolism as well as cell wall homeostasis are important aspects of core stress signaling in both tissues. The complete list of the genes in the SVM gene cores classified in the 4 clusters (blue, yellow, red and green) can be found in the Suppl. File 4.

### Role of ethylene in the SVM gene cores

Because of its key role in a multitude of stresses, and given the presence of ETR2 and ERF2 in the root core, as well as MKK9 – known to play a pivotal role in the activation of MPK6 under ethylene signaling (35) – in the shoot core, the ETH responsiveness of the genes within the SVM gene cores was investigated. To define a robust list of ETH responding genes, we combined the publicly available data of an ETHYLENE INSENSITIVE 3 (EIN3)-ChIP seq analysis (15) with a set of DEGs under early (4h, GSE14247) and late (24h, GSE83573) ETH treatment, forming an ethylene responsiveness database (Suppl. File 5). More than 50% of the genes in the SVM gene core for shoots and roots were highlighted as ETH responsive (Fig. 5A), underpinning the relevance of ETH in both gene cores. Remarkably, the number of ETH-responsive genes increased to 77% and 62.7% in the blue cluster of shoots and roots cores, respectively, further substantiating the central role of ETH in core stress signaling.

**Figure 5.**
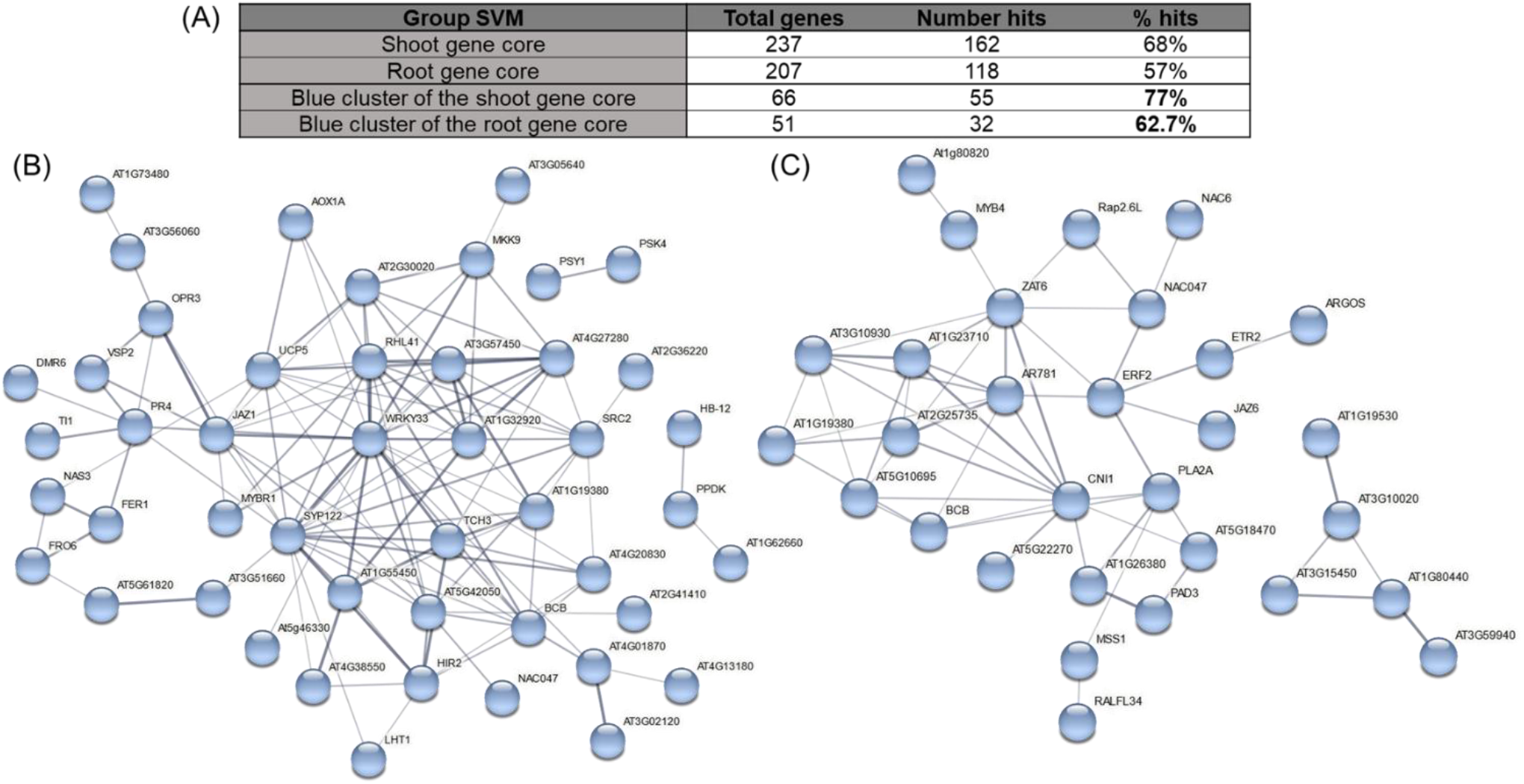
Ethylene-related genes extracted from the shoot and root gene cores. **(A)** Number and percentage of genes of each gene core present in the ETH responsiveness database. Number hits refers to the number of genes from the core present in the ETH responsiveness database (Suppl. File 5), while the percentage indicates the number of hits compared to the total number of genes present in the complete core. **(B and C)** STRING interaction network of the ETH response genes from the blue cluster of the shoot **(B)** and root **(C)** gene cores. The connection between different nodes indicates protein-protein association. Thickness of connecting lines represents the strength of the connections based on experimental data.

The subgroup of genes of the blue clusters detected as ETH-related genes were used to construct a protein interaction network (hereafter denominated as ETH-related clusters). In the case of the shoots, the ETH-related proteins showed the same interconnected pattern as in the complete network (Fig. 3B and 5B). Moreover, WRKY33 still appeared as a central node in shoot stress signaling, together with MKK9. Notably, MKK9, along with MPK3 and MPK6, has been directly linked to both ETH synthesis and signaling (35,36). In addition, LYSINE HISTIDINE TRANSPORTER 1 (LHT1), an amino acid importer shown to be responsible for the transport of the ETH precursor 1-aminocyclopropane-1-carboxylate (ACC; 37) was also part of the ETH-related shoot cluster, as well as the mitochondrial AOX1a which connects the regulation of respiration to stress signaling in an ETH-dependent manner (38).

In roots, the number of genes constituting the interaction network was significantly reduced (Fig. 4B and 5C). Consequently, the number of interactions was also decreased. Nevertheless, the ETR2-ERF2 module again appeared at the center of the interaction network, coordinating other nodes of the network. In addition, the core network also contained AUXIN-REGULATED GENE INVOLVED IN ORGAN SIZE (ARGOS), which is part of a negative feedback mechanism to attenuate ETH responses, further highlighting the importance of coordinated ETH signaling (39). Several TFs that have been experimentally linked to specific stresses or processes, including RAP2.6L/ERF113 (wounding), NAC047 (partial submergence), and NAC6 (leaf senescence), also appeared in this cluster, suggesting a more general function for all.

The high degree of interconnection of the blue clusters of both SVM gene cores along with the multitude of genes related to ethylene responses supports the contention that ETH plays a central role in core stress responses in both shoots and roots.

### Relevant gene families

Certain gene families were identified as crucial players in the SVM gene cores, such as *EXPs*, *AP2/ERF*s, *WRKYs* and *MAPKs*. In addition, novel gene families were also identified by the SVM Clustering algorithm, including *USPs*. To provide a complete and detailed map of the function of these gene families in stress responses, we investigated the transcriptional alterations of all their family members under all conditions in our meta-analysis (Fig. 6; Suppl. File 6). In addition, as a second layer of biovalidation, experimentally validated data about specific members of the selected families corroborated our results (Suppl. Table 2). A few families are highlighted in the next section; further details are presented in the Suppl. Text.

**Figure 6.**
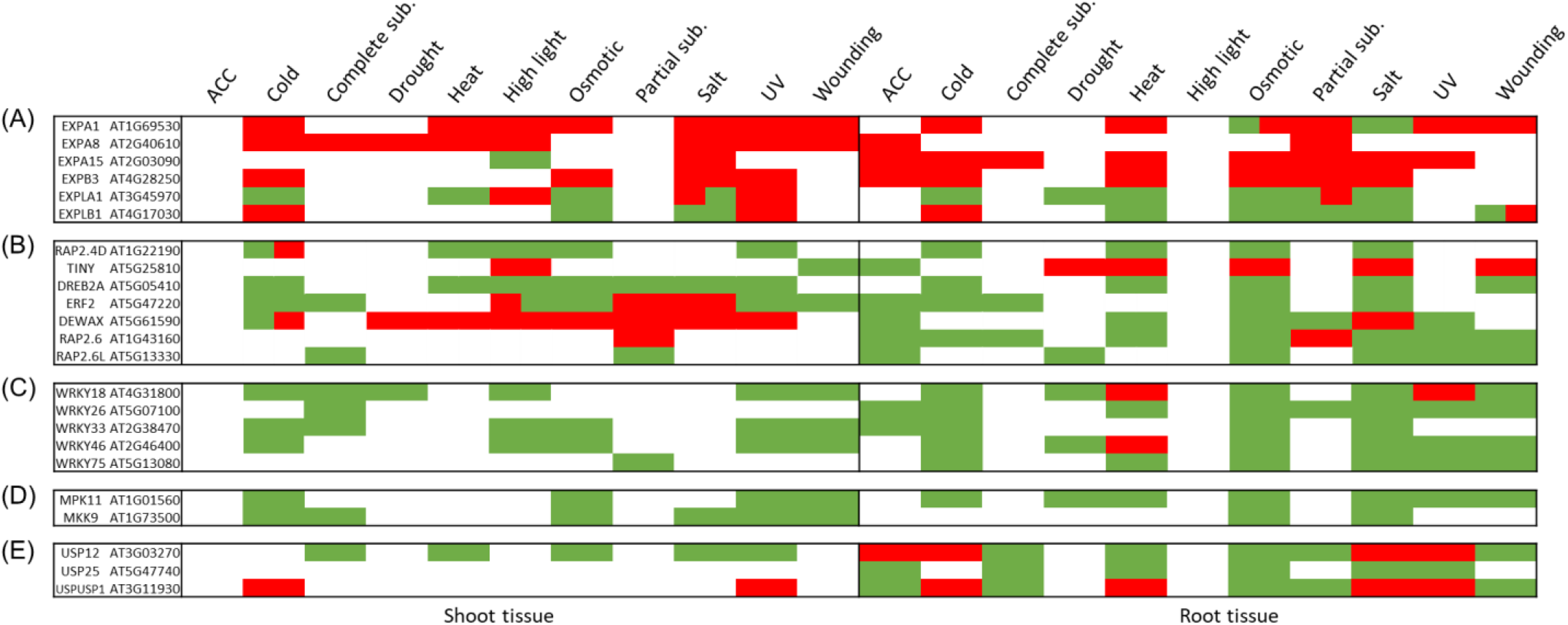
Summary of the presence of different members of expansins (*EXP*) (A), ethylene response factors (*ERF*) (B), *WRKYs* (C), mitogen activated protein kinases (*MAPK*) (D) and universal stress proteins (*USP*) (E) as DEG in the different conditions studied. Green and red color represent up- or down-regulation in each of the conditions. Bicolored cells reflect that the behavior of the gene is variable depending on early or late responses (for complete information on all members of the family, see Suppl. File 6).

## 1. EXPANSIN (EXP) superfamily

*EXPs* were detected as the main enriched gene family in the root core (Fig. S4). They enable cell expansion and increase cell wall flexibility (40,41). The EXP superfamily is divided into four groups: *EXPA*, *EXPB*, expansins-like A (*EXPLA*) and *EXPLB*. In our DEG database, both *EXPA* and *EXPB* subfamilies are predominantly downregulated in several stress conditions with tissue-specific patterns (Fig. 6A; Suppl. File 6). Three members of *EXPA* (*EXPA1*, *EXPA8* and *EXPA15*) and 1 member of *EXPB* (*EXPB3*) were part of SVM stress cores. While transcriptional alterations of *EXPA8*, *EXPA15* and *EXPB3* were observed for roots under certain conditions, *EXPA1* showed transcriptional alterations in all tissues and timepoints in 7 out of 10 studied stresses. These findings highlight the importance of the tissue-specificity of *EXPs* in stress signaling and of *EXPA1* as a main stress regulator. Interestingly, though most *EXPs* were downregulated in response to stress, the *EXPLA* and *EXPLB* groups (with 2 genes present in the SVM gene cores, *EXPLA1* and *EXPLB1*) were upregulated under several stress conditions, suggesting a potential positive role for these subfamilies in stress responses.

## 2. ETHYLENE RESPONSE FACTOR (ERF) family

The ERF TF family is considered to be crucial in both growth and defense responses (42). ERFs are part of the AP2/ERF superfamily, comprising 147 TFs divided in three sub-families: 18 *AP2s*, 122 *ERFs* and 6 *RELATED TO ABSCISIC ACID INSENSITIVE 3/VIVIPAROUS 1* (*RAVs*), as well as a not yet classified gene (AT4G13040) (43). Whereas few to no transcriptional alterations were found for the AP2 family, the RAV and ERF subfamilies revealed to be highly affected by stress, often with very distinct expression patterns (Fig. 6B; Suppl. File 6).

Almost all *ERF* subgroups displayed substantial transcriptional changes under the studied stress conditions (Suppl. File 6). Seven ERFs were classified as members of the SVM gene core, 6 in the root core (*ERF2*, *TINY (ERF040)*, *DREB2A (ERF045), DEWAX* (ERF107), *RAP2*.*6 (ERF108)*, and *RAP2.6L (ERF113)*), and only one (*RAP2.4D (ERF058)*) in the shoot core (Fig. 6B). Interestingly, some of these ERFs showed tissue-specificity. For instance, *RAP2.6/ERF108* and *RAP2.6L/ERF113* showed a similar transcriptional pattern in root tissue, yet exhibiting distinct expression patterns in shoots. While *RAP2.6/ERF108* was downregulated under the early exposure to partial submergence, *RAP2.6L/ERF113* was upregulated under both types of submergence. In contrast, *ERF2* and *DREB2A/ERF045* were broadly expressed in both roots and shoots in most conditions. Since the direct relationship between the members of the ERF family and ethylene is not always clear, we investigated their responsiveness to ACC as well as to ethylene. *ERF2*, *RAP2.6/ERF108* and *RAP2.6L/ERF113* were found to be upregulated by ACC, as well as present in the ethylene responsiveness dataset. The other ERFs were either only upregulated by ACC (*TINY/ERF040* and *DEWAX/ERF107*), only ethylene responsive (*DREB2A/ERF045*), or not detected under ACC, nor ethylene treatment (*RAP2.4D/ERF058*).

## 3. Mitogen activated protein kinases (MAPKs) superfamily

MAPKs are important signaling proteins in many intracellular responses to developmental, physiological and/or environmental stimuli (44). MAPK signaling cascades are typically characterized by a sequence of phosphorylation and activation events along three levels comprising members of Mitogen Activated Protein Kinase Kinase Kinases (MAPKKK, MKKK or MEKK), Mitogen Activated Protein Kinase Kinases (MAPKK or MKK), and Mitogen Activated Protein Kinases (MAPK or MPK). In *A. thaliana* 69 MAPKKKs, 10 MAPKKs and 20 MAPKs have been described (44).

Four out of the ten members of the *MAPKK* group were not responding to any of the conditions studied (Suppl. File 6). However, of the stress-responsive *MAPKKs*, *MKK9* stood out, being upregulated by 6 different conditions, mainly in shoot tissue (Fig. 6D). High salinity and osmotic stress elevated *MKK9* transcription in roots. Not surprisingly, MKK9 appeared as part of the shoot core (Fig. 3B), indicative for its central regulatory role in abiotic stress responses.

Out of 20 *MAPKs,* only 6 did not show a transcriptional effect under any of the stress conditions. Conversely, *MPK11*, *MPK3*, *MPK5* and *MPK19* stood out given transcriptional alteration under 7, 6, 5, and 4 different stress conditions, respectively (Suppl. File 6). However, only *MPK11* was retained by the SVM Clustering as a stress core gene (Fig. 6D). Cold, osmotic and UV stresses upregulated *MPK11* expression in all tissues, while salt induction was root-specific. Interestingly, wounding (shoots and roots), drought (roots) and heat (roots) induced *MPK11* transcription predominantly at early timepoints, implying its relevance specifically during the initial stages of the stress response.

## 4. Universal stress protein (USP) superfamily

USPs are proteins involved in, as their name suggests, a broad range of metabolic processes related to biotic and abiotic stress, such as nutrient starvation, heat shock and oxidative stress (45). Nevertheless, their specific roles and molecular mechanisms remain largely unknown. In *A. thaliana*, 41 genes encode for USP proteins, cataloged according to domain organization. From our analysis, the *USP* family and the single gene belonging to the double USP domain group (*USPUSP*) appear to be involved in all stress conditions (Suppl. File 6).

USP12, USP25 and USPUSP1 were detected as part of the root gene core. *USP12*, characterized as a gene involved in ROS modulation in anoxia conditions, is upregulated in submergence conditions, but also in heat and osmotic conditions in both tissues (Fig. 8E; Suppl. Table 2). *USP25* and *USPUSP1* appear to be tissue-specific players in the general stress response, being only expressed in roots. It is evident that the function of USPs deserves more scrutiny, given their highly specific expression patterns as well as the central role of certain family members in general stress signaling.

### Experimental validation: the role of EXPAs, MKK9, and LHT1 in general stress responses

As part of a third layer of biovalidation supporting the role of members of the gene cores as central stress regulators, we first studied the transcriptional alterations of 3 members of the EXPA family (*EXPA1*, *EXPA10* and *EXPA14*), represented in the root core, and EXPA1 as a part of both gene cores, using translational reporter lines (*pEXPA1::EXPA1-mCherry* (Fig. 7A), *pEXPA10::EXPA10-mCherry* (Fig. 7B), *pEXPA14::EXPA14-mCherry* (Fig. 7C) (40). We exposed the three lines to cold, salt and osmotic stress for 1 hour. The expression patterns in control conditions corresponded with the patterns described by Samalova *et al*. (2023) (40). Short-term exposure to stress induced changes in the intensity levels, indicating a change in the quantity of EXPA present in the studied tissue. Specifically, salt treatment significantly increased the levels of EXPA1 and EXPA10, while decreasing the level of EXPA14 (Fig. 7D), corroborating the expression changes after the equivalent treatment obtained in our meta-analysis (Fig. 7E). Osmotic treatment modestly increased the levels of EXPA1 and EXPA10, without affecting the level of EXPA14. Upon cold treatment, none of the levels of the studied EXPAs differed from those of the control samples, again confirming the expression data. In conclusion, the data of the translational reporter lines matched with the transcriptional data derived from the meta-analysis, experimentally validating the robustness of our analysis, and positioning EXPAs as key regulators of multiple stress responses as evidenced by their relevance in both shoot and root gene cores.

**Figure 7.**
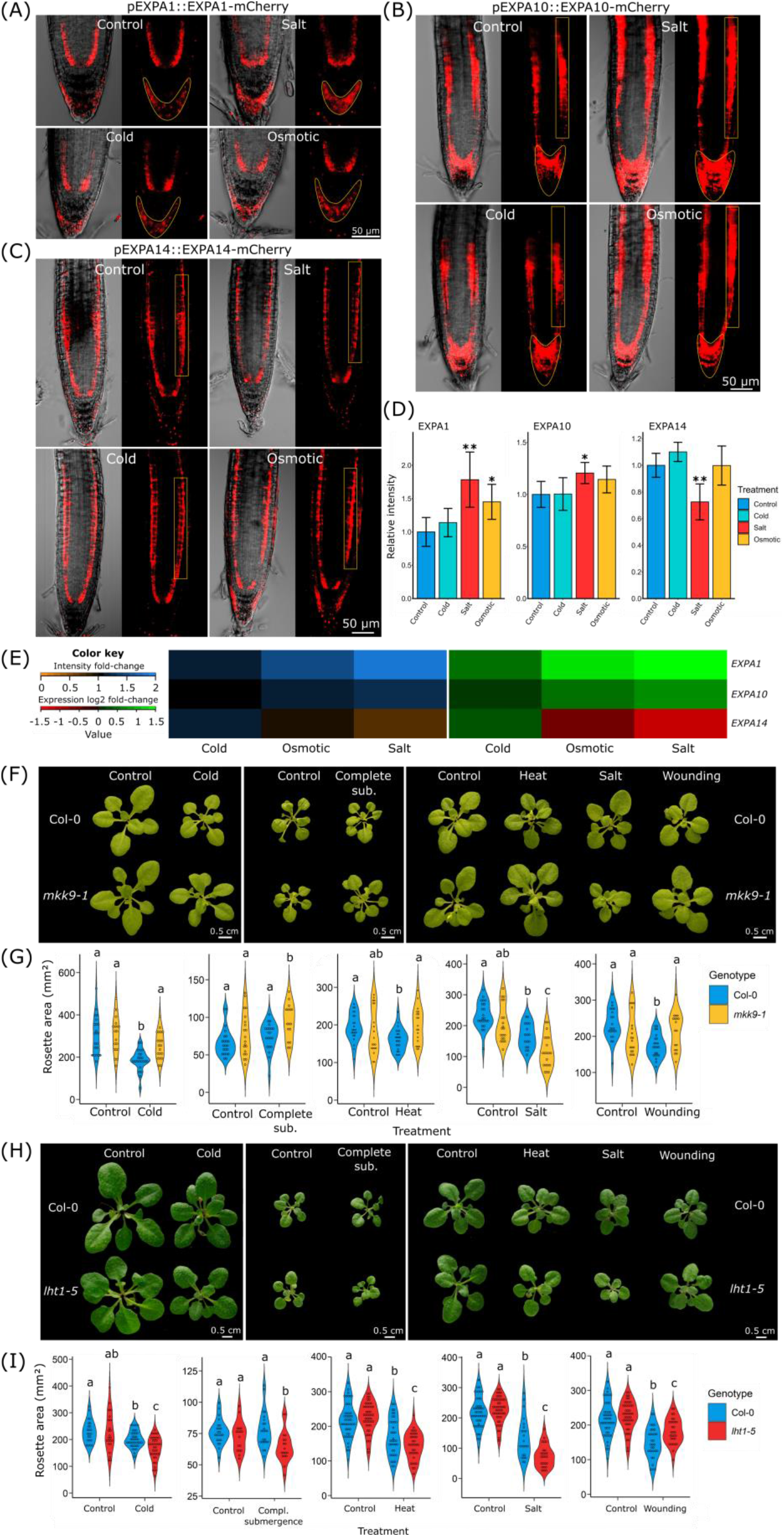
Experimental validation of the transcriptional alterations of EXPAs and the role of MKK9 and LHT1 as central hub in stress responses. Fluorescence intensity of the translational reporter *lines pEXPAx::EXPAx-mCherry* for EXPA1 **(A)**, EXPA10 **(B)** and EXPA14 **(C)** was measured in the lateral root cap for EXPA1, in the root tip and a region of the cortex for EXPA10 and in a region of the cortex for EXPA14. The measured region of interest (ROI) is highlighted by a yellow box. Scale = 50 µm. **(D)** Fluorescence intensity changes in relation to control conditions (n > 15 seedlings per sample). A *p*-value of *p* < 0.05 was assumed as statistically significant (* < 0.05; ** < 0.01; *** < 0.001). **(E)** Heatmap showing intensity fold-changes represented as relative marker gene intensity compared to control conditions (ranging from 0 (orange) to 2 (blue)) and expression fold-changes based on transcriptomic analyses (represented as the log2-fold change compared to control conditions, ranging from -1.5 (red) to 1.5 (green)) for each stress condition. **(F-I)** Rosette area of Col-0 and *mkk9-1* (F-G) *lht1-5* (H-I) plantlets. Rosette area was measured after 4 days of cold (4oC), complete submergence, heat (42oC), salt stress (100 mM NaCl), and wounding, and after 10 days in recovery conditions (n > 20 plants per sample). Scale = 0.5 cm. A *p*-value < 0.05 was assumed as statistically significant and different letters represent statistical differences between samples.Shapiro’s test and Levene’s test were used to assess normality and variance of each dataset. To compare the intensity of the different treatments with the control samples, Student’s T-test and Wilcoxon rank sum exact test were used for parametric and non-parametric testing, respectively.). To compare rosette areas, a two-way ANOVA followed by post-hoc Tukey’s test were used as parametric tests, while a Kruskal-Wallis rank sum test followed by Dunn’s Multiple Comparison test were used as non-parametric tests..

Secondly, we investigated the function of MKK9, as one of the main proposed regulators in the shoot core (Fig. 3B). We assessed stress tolerance by comparing alterations in rosette growth between the loss-of-function mutant *mkk9-1* and the wild-type Col-0 (Fig. 7F-G). To consider a representative set of stress conditions, we selected 5 of the 10 conditions studied in our analysis, covering each of the clusters obtained by HCA, namely complete submergence, cold, heat, salt and wounding (Fig. 1).

As observed in Fig. 7, *mkk9*-*1* mutants behaved differently from the wild-type in all tested conditions. In the wild type, cold, heat and wounding led to a significant reduction in rosette area in comparison with non-treated samples (Fig. 7F-G). In contrast, *mkk9-1* growth was not compromised by these treatments and rosettes were significantly larger than those of treated Col-0 plants, showing an increased tolerance of the mutant. Furthermore, after salt treatment, Col-0 showed a decrease in rosette size, while *mkk9-1*, in contrast to the other conditions, was hypersensitive to salinity with an even larger reduction in rosette size. Finally, when the plantlets were maintained for 4 days in submerged conditions, rosette growth was positively affected in *mkk9-1* plantlets, with rosettes being significantly larger compared to both the controls and treated wild-type.

Lastly, we analyzed the response of the *lht1-5* loss-of-function mutant to the different stress treatments (Fig. 7H-I). Both its presence as a member of the shoot core as well as its function in amino acid and ACC transport suggest that LHT1 could act as a vital component of general stress signaling. Indeed, plantlets that lack functional LHT1 are hypersensitive to cold, complete submergence, heat, and salt stress. In contrast, *lht1-5* plantlets were less affected than Col-0 plants upon wounding.

The results of the three above-mentioned analyses empirically corroborate the suitability, robustness and reproducibility of the SVM gene cores and, at the same time, further validating the general role of these genes in the regulation of stress responses in plants.

## Discussion

### Machine learning as a tool for rapid identification of the stress gene core: strengths and limitations

Machine learning approaches represent a powerful tool for data analysis. Yet, the design, scientific question and biological significance need to be carefully addressed in order to avoid incorrect interpretations or lack of reproducibility of the generated output (26). For that reason and in addition to the robust design of our computational analysis, we have included a three-layered biovalidation process. Firstly, we have used experimentally determined stress markers to assess the quality of the generated DEG libraries (Suppl. Table 1). Secondly, expression patterns of specific members of gene families with well-characterized transcriptional behavior under different stress conditions were used as an additional level of validation (Suppl. Table 2). Finally, in order to validate the genes forming the proposed SVM gene core, we empirically analyzed the effect of stress exposure on alterations in transcription (Fig. 7A-D) and the function of three key genes in physiologically representative stress conditions, one of which was previously not linked to abiotic stress conditions (Fig. 7F-I). Altogether, this provides a solid basis to put confidence in the previously uncharacterized genes which are part of the stress gene core, including *USPs*, offering strong evidence towards their putative biological function. In addition, it strengthens the validity of our methodology to study complex processes. The proof of concept presented in this study could be extended to, for example, the determination of the central players in biotic stress responses, as well as to deeper understanding of the differences between the responses induced by necrotrophic, biotrophic, and hemibiotrophic pathogens.

### To stress stimuli and beyond: the role of the SVM gene cores in stress responses

Most environmental changes are first sensed by displacement of the cell wall-membrane interface (31). Plant genomes have evolved specific mechanisms to monitor and ensure membrane integrity and cell wall rigidity (46,47). Representatives of these mechanisms were found in the SVM gene cores. *EXPANSINs*, a gene family extensively studied for its implication in cell wall loosening and cell growth (40), was the main family enriched in the root core (Fig. S4) and, moreover, *EXPA1* was included in the 19 genes shared between root and shoot cores (Fig. 6A). This supports the contention that cell wall loosening and remodeling, apart from its role in normal growth, is also crucial for the adaptation to environmental stresses, especially in roots. Typically, stresses that lead to ROS production and loss of water alter the expression of EXPs (41). However, the precise mechanism of action of EXPs in stress mitigation remains unclear. To provide empirical validation of our meta-analysis as a third biovalidation layer, we compared the effects of short-term stress exposure on EXPA1, EXPA10 and EXPA14 accumulation with the transcriptional changes detected in our analysis revealing a robust parallelism between the two datasets (Fig. 7A-E). *EXPA* and *EXPB* are well-characterized groups and were downregulated in the majority of stress responses (Fig. 6A, Suppl. File 6), supporting the importance of cell wall and membrane rigidity in stress responses. Among *EXPAs* and *EXPBs*, *EXPA1* stands out as a pivotal gene in stress responses, while the other members seems to have a more specific role. The other two *EXP* groups, and especially *EXPLA1* and *EXPLB1* showed an opposite trend, being mainly upregulated. Since the functions of both subgroups have not been elucidated up to date and our data suggest that they could play opposite roles compared to *EXPA* and *EXPB*, it will be interesting to further unravel their roles.

Alterations at the level of the cell membrane often serve as initiators of stress signaling, with the activation of membrane-anchored Ca^2+^ channels as one of most relevant stress response inducers (48). Calcium influx and signaling have been implicated in drought, cold, salt, osmotic, hypoxia and flooding responses, highlighting their importance in most stresses (49,50). Nevertheless, little is known about the specific genes controlling this signaling network (57). One of the gene families in the shoot core corresponded to ion exchangers, specifically cation/Ca^2+^ exchangers (Fig. S4). *CALCIUM EXCHANGER 1* (*CCX1*), upregulated in UV, complete submergence, wounding, osmotic and salt stresses during the early exposure to stress in shoots. On the other hand, its paralog *CCX2* is present in the root gene core, upregulated in drought, cold, heat, osmotic and salt stresses, also during the early exposure. Both were previously linked to ROS accumulation, while a *CCX2* loss-of-function mutant displayed hypersensitivity to salt stress (51,52). The presence of *CCX1* and *CCX2* in the shoot and root cores, respectively, highlights their potential involvement during the early stress responses. In addition, they could be excellent targets for the study of Ca^2+^ channels in the systemic communication between root and shoot.

In addition to Ca^2+^ influxes, interpretation of Ca^2+^ waves by Calmodulins (CaM) and calmodulin-like (CML) proteins is required for adequate stress signaling and inter-organ communication (53). Recently, the role of *TCH2/CML24* has been demonstrated to be vital for heavy metal tolerance in *A. thaliana* by its interaction with WRKY46 (present in the shoot gene core; Fig. 5A) (54). Another CML, TCH3/CML12, plays a central role in the interaction network of the shoot core, interacting with the central protein WRKY33. Moreover, the fact that these genes have been related to other stresses apart from the ones included in our analysis (such as heavy metal tolerance), endorses the general nature of this core in stress signaling. On the other hand, it can verify genes that are playing a central role in the interaction network (*TCH3-WRKY33*) as key players in general stress response coordination.

Following the trip of Ca^2+^ waves, MAPK signaling cascades are highlighted as one of the main coordinators, facilitating downstream processes (44). Nevertheless, the study of MAPKs is hindered by the complexity of MAPK signaling cascades and their regulation (55). Certain MAPKs arose in our analysis, such as MKK9, whereas others, such as MPK6 – shown to be part of the senescence-related module MKK9-MPK6 but mostly controlled by post-translational regulation (55) – did not appear. Our study revealed an interesting and particular role of MKK9 in the shoot interaction network, binding to the central WRKY33 and TCH3 (Fig. 3B). Functional analysis of a loss-of-function *mkk9-1* mutant supported its role in the coordination of stress responses (Fig. 7F-G). In addition, the results corroborated previous observations, where the *mkk9-1* mutant exhibited hypersensitivity to high salinity (36). This biovalidation not only supports the potential role of the SVM gene cores in stress responses, but also provides strong evidence for the role of MKK9 as a part of a hub in stress responses, giving additional insights into its biological function.

In many stress signaling cascades hormones are activated as key mediators after initial stress sensing and signal relay. Ethylene is such a key stress hormone, with a negative effect in cold tolerance (56). Conversely, ethylene positively influences the survival rate and tolerance to flooding conditions, mediated by ERF-VIIs (57), and it also improves salt tolerance through *ERF1* induction (58). This study provides evidence for broad transcriptional alterations of *ERFs* in multiple stress conditions. Especially *ERF2*, upregulated in 5 and 6 conditions in roots and shoots respectively, came forward as a central player in the root core interaction network (Fig. 6B). In addition, the ERF2–ETR2 module demonstrated that one of the main root core functions is related to ethylene responses, supporting the pivotal role of ethylene in root stress responses (Fig. 4F). Interestingly, the amino acid transporter LHT1 was part of the shoot stress core (Fig. 3B). LHT1 was previously shown to transport ACC in *Arabidopsis*, and *lht1-5* mutants display an early-senescence phenotype (37). Here, we show that loss of *LHT1* leads to an altered tolerance to all of the tested stresses (Fig. 7H-I). These results indicate that LHT1 plays a prime role in abiotic stress responses in addition to its previously reported function during pathogen infection (59). Though LHT1 clearly appears to act as an important node that simultaneously regulates cellular ACC availability – and thus ETH – as well as levels of other (non)-proteinogenic amino acids, more work on its precise mode of action is needed. Besides this direct link, using our ethylene-responsiveness database we were able to detect the relevance of ethylene in the regulation of more than the 50% of the genes in both gene cores (Fig. 5). In addition, some of the abovementioned key genes (*CCX1, TCH3, WRKY33* and *MKK9*) were also related to ethylene responses, further supporting the pivotal role of ethylene in expression of these central core genes (see Suppl. Fig. 5 for a schematic overview of the process).

### Conclusions

In conclusion, we demonstrated the suitability of an unsupervised machine learning technique, SVM Clustering, to gain insights in complex biological processes. The proposed methodology was proven robust and provides a comprehensive view on plant responses to a particular set of growth conditions. The stress cores cover a number of genes previously linked to specific stresses, corroborating the solidity of the cores and emphasizing the power of the SVM Clustering approach to identify genes involved in general stress signaling, a stepstone for studies on impact of global climate change on plants. Moreover, the approach allows to uncover novel genes as central players in the processes under study, with either another unrelated or no function previously assigned. Hence, SVM Clustering is a robust mining tool to rapidly gain a holistic view on a gene network at the center of set of responses. Furthermore, we have demonstrated the vital role of ETH signaling in the core stress signaling network. Last but not least, during the analysis, different databases were generated unifying valuable information regarding plant stress-responsiveness. Those datasets present comprehensive expression patterns of complete stress-related gene families (*EXP*, *WRKY*, *AP2*/*ERF* and *MAPK*) as well as the previously poorly characterized *USPs* and the *EXP* subgroups *EXPLA* and *EXPLB*, providing new insights into their biological roles. These will nurture future functional analyses of yet uncharacterized genes and relevant members within large gene families, unchartered territory that hampers full understanding of complex stress responses in plants.

While corroborating the role of a number of genes known to be stress-related, the gene cores present excellent novel candidates for the engineering of plant tolerance to a wide range of adverse conditions, surpassing the limitations of single stress-related empirical studies including single transcriptomic analyses. Overall, the developed methodology, along with the derived datasets, represents a stepstone towards future sustainable agriculture.

## Materials and methods

### Database selection

*A. thaliana* transcriptomes were obtained from the Gene Expression Omnibus (GEO) database (https://www.ncbi.nlm.nih.gov/geo/; Accessed in June 2021). The keywords “Arabidopsis” and “stress” provided a list of 945 series of data after filtering for organism (“*Arabidopsis thaliana”*; filter aimed to exclude analyses using *A. thaliana* genes in other species) comprising both microarrays and RNA-seq platforms. Keywords for specific conditions were used to refine the search (for more details, see Suppl. text). For each stress type, the datasets corresponding to specific time points were selected, taking the following criteria into account: (1) at least duplicates per treatment, (2) availability of raw data, (3) tissue specificity of raw data is known (root or shoot), and (4) the age of analyzed plants was between 1 and 4 weeks, reducing the bias of the developmental stage. The final list was composed of 23 data series comprising 500 single transcriptomes (Suppl. File 1). The sets were divided according to temporal and spatial determinants, with different time points divided into early (1, 3 and 6 hours) and late responses (12 and 24 hours), and tissue types (root and shoot tissues) (Suppl. Table 1). The selection of time points is related to the variability of differentially expressed genes (DEG) peaks in literature, the maximum usually being between 1 and 3 hours for early responses but displaced in some examples up to 6 h (60,61). In this way, the inclusion of a 6 hours timepoint in the category of early responses encompasses most early transcriptomic changes. Only the datasets containing information for root early and late and/or shoot early and late responses were considered for subsequent analysis, thus excluding cadmium and hypoxia. All datasets used are specified in the Suppl. File 1.

### Data pre-processing, normalization, and differentially expressed genes (DEGs) detection

Data pre-processing and normalization were performed using the Robust Multi-array Average (RMA) method, considered the best method to increase comparability between different platforms (62). Unexpressed tags were detected and filtered by the MAS 5.0 method (tags with an alpha value > 0.06 were discarded, as default parameter defined by Affimetrix) and by using the *edgeR* package, for microarray and RNAseq respectively, and redundant tags were expressed as the average of the redundant probes. All expression data were expressed as log2/log2-CPM to increase comparability. Low quality datasets were removed from pre-processed data (63). DEGs in each individual dataset were detected by computing a linear model using an empirical Bayes method, which has been demonstrated to provide high-precision analyses of transcriptomic data in both microarray and RNA-seq analyses (62). Genes with a log-fold change (LFC) > |1| (equivalent to a fold change > |2|) and a false discovery rate (FDR; obtained by the Benjamini and Hochberg corrected *p*-value (64)) < 0.05 were selected as DEGs. This strict threshold was defined in order to provide a DEG list containing only the most relevant up- or down-regulated genes.

### Hierarchical clustering analysis (HCA) of the different stress modules

Hierarchical clustering analysis (HCA) was selected as an unsupervised machine learning method to detect similarities and differences between the DEGs modules of each stress, taking into account time and tissue. DEGs modules were created considering up- and down-regulated DEGs in each stress condition for early (1, 3 and 6 hours) and late (12 and 24 hours) time points for both root and shoot tissues. In case of discordance between some of the points (when a gene was up-regulated in one timepoint but down-regulated in other, or *vice versa*) the earliest time point for the early module and the latest time point for the late module were selected as representatives. The computation of the clusters was performed by an agglomerative HCA method with the Gower’s distance as a dissimilarity matrix (65). To assess the differences between the four computed hierarchical clusters, pairs of dendrograms were compared and differences were quantified by the Baker’s gamma index (for a detailed overview of the process see Suppl. text).

### Meta-p-value computation and Support Vector Machine (SVM) Clustering based classification

To overcome the variability of sample-specific transcriptomes, the meta-*p*-values for each stress, tissue and timepoint were computed by the generalized weighted Fisher’s method with sample-sizes correction (66). The selection of this method is based on the meta-analysis decision scheme proposed in (24), considering the source data (different platforms) and the heterogeneity of the dataset. The Benjamini and Hochberg corrected *p*-values obtained from the individual DEG detection analysis were used as input (67). As output, a complex dataset composed of 11 meta-*p*-values (one per stress condition and ACC treatment), per tissue and timepoint for each gene is given. Due to the multidimensionality of the dataset, we used SVM Clustering as an unsupervised classification method for the definition of the ‘stress gene cores’ (28). SVM Clustering takes advantage of all the machinery provided by standard SVM to create a partition in a dataset, i.e., compute the decision boundaries between a set of user pre-defined clusters. In addition, since SVM Clustering is an unsupervised machine learning method, it significantly reduces the potential bias inherent to supervised learning methods, as it does not depend on previous knowledge of the data. We selected the Radial Basis Method (RBF) as SVM kernel function, highly powerful for non-linear high-dimensional datasets and allowing for a more accurate classification (68). The parameters cost (*c*) and gamma (*γ*), crucial for robust and trustable results, together with the chosen kernel method, were optimized individually for each tissue by time-set (root-early, root-late, shoot-early, and shoot-late) using grid-search (69). The output consisted of four stress gene cores, denoted as SVM gene cores.

### Gene core analysis

The comparisons between the SVM gene cores were performed by computing the overlap between different datasets and were visualized using Venn Diagrams (*VennDiagram* R package was used). The hypergeometric test was used to assess the significance and over- or underrepresentation of the different overlapping groups of genes as described in (70) (for more details see Supplementary Text).

For GO enrichment, the *topGO*, *goProfiles* and *clusterProfiler* R packages were used.

GenFAM (Gene Families) (71) and STRING (Search Tool of Recurring Instances of Neighbouring Genes; accessed in August 2022; version 11.1) (72) were used for gene family enrichment and the interaction network construction, respectively. For SVM gene cores clustering, the *k*-means method from the STRING online tool was used.

### Detection of gene responsiveness to ethylene

To detect the relation between the SVM gene cores and ETH responses, experimentally validated EIN3 interactions (15) and the transcriptomic response to ETH were used (GEO accession numbers: GSE14247, GSE83573) to create an ETH responsiveness database (Suppl. File 5). The subset of ETH responsive genes within the SVM gene cores was used as input in STRING (version 11.1) (72), and their interaction network was obtained.

### Plant material and analysis of stress effects

The analysis of EXPA levels was performed using the *pEXPAx::EXPAx-mCherry* translational reporter fusions for the genes *EXPA1*, *EXPA10* and *EXPA14* (42). Seeds were surface-sterilized according to (73) and subsequently plated on half-strength Murashige and Skoog medium containing 1% sucrose and 0.8% agar (hereafter MS/2). After 3 days of stratification at 4°C, plates were transferred to a tissue culture room and grown in a 16/8-h photoperiod (70 µmol ⋅ m^−2^ ⋅ s^−1^) for 4 days at 21°C, placed in a vertical position. Subsequently, the plantlets were exposed to a short-term (1 hour) stress treatment (for more details see Supplementary Text). Confocal laser scanning microscopic images were obtained with an inverted Nikon TiE-C2 confocal microscope. Roots were imaged with a 20x CFI Plan Apochromat VC objective lens (NA 0.75, dry). Images (1024 x 1024; 1.5x scanner zoom) were collected by exciting mCherry with a solid-state 561 nm laser and emission was collected from 571 to 700 nm. The same settings were kept to compare fluorescence intensities between treatments of specific transgenic lines. Quantification of the mCherry signal was performed in ImageJ. Regions of interest (ROI) were defined for each line focusing on the expression patterns as described by Samalova et al. (2023). Signal intensity was quantified as gray value intensity normalized by the ROI size in µm^2^. Shapiro’s test and Levene’s test were used to assess normality and variance of each dataset. To compare the intensity of the different treatments with the control samples, Student’s T-test and Wilcoxon rank sum exact test were used for parametric and non-parametric testing, respectively. A p-value of p < 0.05 was assumed as statistically significant (* < 0.05; ** < 0.01; *** < 0.001).

For the analyses of *lht1-5* and *mkk9-1* mutants, *A. thaliana* Columbia (Col-0) was used as control. The *mkk9-1* and *lht1-5* mutants (in Col background) was obtained from the NASC Arabidopsis stock center (SALK_017378 and SALK_115555C, respectively). Seeds were surface-sterilized, plated on MS/2 medium, stratified and transferred to a tissue culture in the same conditions as mentioned above. Hereafter, plantlets were exposed to their respective long-term stress treatments (for more details see Supplementary Text). After 10 days of recovery, plants were imaged (CANON EOS 550D camera (Canon, Tokyo, Japan)) and the rosette area was analyzed using the ImageJ plug-in (National Institutes of Health) of *Rosette tracker* (74). Rosette areas were plotted as violin plots using the *ggplot2* R package. To compare rosette areas, a two-way ANOVA followed by post-hoc Tukey’s test were used as parametric tests, while a Kruskal-Wallis rank sum test followed by Dunn’s Multiple Comparison test were used as non-parametric tests. A p-value < 0.05 was assumed as statistically significant and different letters represent statistical differences between samples.

## Acknowledgements

DVDS was responsible for the design and supervision of the biological aspects of this work, while IZ was responsible for the design and supervision of the computational aspects. RSM performed the datamining, the meta-analysis and the machine learning approach. RSM, together with TD, performed the functional validation of MKK9 and LHT1. MS and JH provided the EXPA translational lines and guided their analysis. TD performed the EXPA translational reporter lines analysis. RSM, TD, IZ and DVDS prepared the manuscript. All authors reviewed the manuscript.

This work was supported by grants from Ghent University (Bijzonder Onderzoeksfonds BOF-BAS) and the Research Foundation Flanders (FWO; G032717N and G082421N) to DVDS. IZ thanks the program ‘*Maria Zambrano para la atracción del talento internactional*’ (ref. 2022UPC-MZC-94008) from the Ministerio de Universidades (Spain). RSM is grateful to FWO (grant number 1288923N) for a senior postdoctoral fellowship.

The authors declare that they have no competing interests. All data needed to evaluate the conclusions in the paper are present in the paper.

Supplementary information will be available in the final published version.

